# Deletion of the ion channel *Trpm4* increases cardiac inflammatory markers and fibrosis after myocardial infarction in mice

**DOI:** 10.1101/2022.10.24.513479

**Authors:** Mey Boukenna, Jean-Sébastien Rougier, Parisa Aghagolzadeh, Sylvain Pradervand, Sabrina Guichard, Anne-Flore Hämmerli, Thierry Pedrazzini, Hugues Abriel

## Abstract

**BACKGROUND:** The first cause of mortality worldwide is ischemic heart disease. In myocardial infarction (MI), the ischemic event causes cell death, which triggers a large inflammatory response responsible for removing necrotic material and inducing tissue repair. Endothelial cells, immune cells and fibroblasts play a key role in orchestrating this healing process. TRPM4 is a Ca^2+^-activated ion channel permeable to monovalent cations and its silencing or knocking out was shown to critically modify cellular functions of these non-myocytic cell types.

**OBJECTIVE:** Our aims were to 1) evaluate the role of TRPM4 on mice survival and cardiac function after MI; and 2) investigate the role of TRPM4 in the post-MI acute and chronic inflammatory response.

**METHODS:** We performed ligation of the left anterior descending coronary artery or sham intervention on 154 *Trpm4* WT or KO male mice and monitored survival for up to 5 weeks as well as cardiac function using echocardiography at 72h and five weeks. We drew blood at different acute time points (6h, 12h, 24h) and performed time-of-flight mass spectrometry to analyze the sera proteomes. Further, we sacrificed sub-groups of mice at 24h and 72h after surgery and performed single-cell RNA sequencing on the non-myocytic cells. Lastly, we assessed fibrosis and angiogenesis at five weeks using type I collagen and CD31 immunostaining respectively.

**RESULTS:** We observed no significant differences in survival or cardiac function post-MI between both genotypes. However, our serum proteomics data showed significantly decreased tissue injury markers such as creatine kinase M and VE-Cadherin in KO compared to WT 12h after MI. On the other hand, inflammation characterized by serum amyloid P component in the serum, as well as higher number of recruited granulocytes, M1 macrophages, M1 monocytes, Mac-6 macrophages, and expression of pro-inflammatory genes such as *Il1b, Lyz2* and *S100a8/a9* was significantly higher in endothelial cells, macrophages and fibroblasts of KO than of WT. This correlated with increased cardiac fibrosis and angiogenesis 5 weeks after MI in KO.

**CONCLUSION:** Our data suggest that knocking out *Trpm4* drastically increases acute inflammation post-MI, is associated with increased chronic fibrosis and does not improve survival at 5 weeks post-MI. Thus, targeting TRPM4 in the context of MI should be pondered carefully and approaches that nuance the timing of the inhibition or cellular target may be required.

## INTRODUCTION

Ischemic heart disease is a leading cause of death affecting about 126 million patients worldwide ^1^. A majority of these cases is due to chronic atherosclerosis reducing coronary perfusion towards the cardiac muscle. This process can go undiagnosed for decades, until suddenly a triggering event like intense exercise unmasks a mismatch between increased oxygen demand and decreased supply leading to tissue ischemia. When the ischemia lasts longer than a few minutes, cells experience hypoxia and start dying, resulting in myocardial infarction (MI) that is associated with an acute mortality of 34-43%^2^. Unfortunately, a considerable proportion of patients that survive MI develop heart failure, which is mainly driven by adverse cardiac remodeling and cardiac fibrosis. Heart failure is associated with again high mortality and low quality of life despite the available pharmacological treatments^3^. Thus, the need for novel therapeutic approaches post-MI is dire.

In recent years, various studies have identified the ion channel TRPM4 as a potential pharmacological target in myocardial infarction^4–6^. TRPM4 is a calcium-activated ion channel permeable to monovalent cations such as Na^+^, K^+^, Li^+^ or Cs^+ 7^. Its role in cardiac conduction is actively investigated, as many arrhythmic disorders have been linked to mutations leading to gain- or loss of function of the *TRPM4* gene^8–11^. Upon increase in intracellular Ca^2+^, Ca^2+^ binds to TRPM4 inducing a conformational change that opens the channel allowing monovalent cations to move across the plasma membrane. Due to the intra- and extracellular fluid composition, this usually results in a Na^+^-influx into the cell, thereby depolarizing the plasma membrane. In addition to the direct effect of Na^+^ on the action potential in cardiomyocytes ^12–16^, and on endothelial cell death via oncosis ^16^, it also exerts an indirect function by decreasing the driving force for Ca^2+^ ions to enter the cell through Ca^2+^-channels. Since Ca^2+^ is an important second messenger, this indirect effect of TRPM4 on its homeostasis has been shown to have functional implications on cardiomyocyte hypertrophy, macrophage phagocytosis, lymphocyte cytokine production as well as fibroblast migration and contractility^13,17–19^.

Even though the heart is made up of only approx. 30% of cardiomyocytes and approx. 70% of other cells, such as endothelial cells, fibroblasts, immune cells and smooth muscle cells, all previous studies investigating the effect of TRPM4 in MI focused on cardiomyocytes. Previous findings suggest that knocking out *Trpm4* 1) protects against ischemia-associated disturbance of Ca^2+^ homeostasis, and thus reduces the occurrence of lethal ventricular arrythmias and 2) increases Ca^2+^-dependent contractility, improving inotropy upon ß-adrenergic stimulation^13,20^. However, findings on how TRPM4 affects survival and heart function diverge and the role of TRPM4 in MI is still poorly understood. Importantly, endothelial cells are the first key players in the inflammatory signaling pathway, inducing infiltration of immune cells into the injured tissue and complement activation. The subsequent inflammatory response mediated mainly by granulocytes, macrophages and fibroblasts that transdifferentiate into myofibroblasts, is crucial for scar tissue formation and thus recovery of cardiac function^21–23^. The presence of TRPM4 in these cells prompted us to look beyond cardiomyocytes and investigate the role of TRPM4 in the inflammatory response post-MI. Here, we therefore present the first detailed characterization of the acute inflammatory processes of *Trpm4* KO compared to WT mice after MI using single-cell RNA sequencing and proteomic approaches. Briefly, we performed LAD ligation or sham intervention on *Trpm4* WT and KO mice and sampled blood or cardiac non-myocytic cells at early timepoints up to 24h and 72 hours, respectively. We observed decreased levels of tissue-injury markers such as creatine kinase M (CKM) or VE-Cadherin (CDH5) in KO sera compared to those of WT, whereas inflammatory marker serum amyloid P component (APCS) as well as pro-inflammatory cell types were upregulated in KO. This increased inflammation in KO correlated with increased fibrosis and angiogenesis at 5 weeks after surgery as measured by immunohistochemistry suggesting that deletion of *Trpm4* may have deleterious effects on remodelling post-MI.

## MATERIAL AND METHODS

| Product | Referennce | Purchased from company |
| --- | --- | --- |
| Harris hematoxylin | ref# HHS16 | Sigma Aldrich |
| AEC staining kit | ref# ab64252 | Abcam |
| Vecta Stain ABC kit | ref# PK7100 | Adipogen |
| Anti-Collagen-I-antibody | ref# ab21286 | Abcam |
| Anti-SMA-antibody | ref# ab5694 | Abcam |
| Anti-Rabbit-Biotin | ref# ab7089 | Abcam |
| Anti-Rat-Biotin | ref# 559286 | BD Biosciences |
| Cardiomyocyte isolation kit | ref#88281 | Thermofischer |
| Calcein-AM | ref#564061 | BD Biosciences |
| DAPI | ref# 564907 | BD Biosciences |
| NovaSeq 6000 S1 Reagent Kit v1.5 | ref#20028317 | Illumina |

### Mice

8-12 weeks old male mice were used in all experiments. We either used B6.Cg-Trpm4tm1.2-PG or C57Bl/6N wild-type mice from Janvier Labs. We engineered the *Trpm4* ^-/-^ (B6.Cg-Trpm4tm1.2-PG) mouse line, which is a constitutive global knockdown of the *Trpm4* gene, by excision of the exon 10 and backcrossing was performed using C57Bl/6N wild-type mice from Janvier Labs. All mice were accommodated to the vivarium of the experimental facility for at least two weeks prior to any experimental procedure. Mice were kept in cages with a maximum of five mice per cage and with unrestricted access to food and water. All genotypes were controlled again using PCR at the end of the study. Mouse experiments have been approved by the Government Veterinary Office (Lausanne, Switzerland) and conducted according to the University of Lausanne institutional guidelines and the Swiss laws for animal protection as well as the guidelines from Directive 2010/63/EU of the European Parliament on the protection of animals used for scientific purposes. Accordingly, all euthanasia was performed by neck dislocation after anesthesia by an intraperitoneal injection of a mixture of ketamine/xylazine (65 and 15 mg/kg respectively), i.e. an injection volume of 0.1 mL per 10g of body weight. The overall experimental setup is visualized in Figure 1.

**Figure 1.**
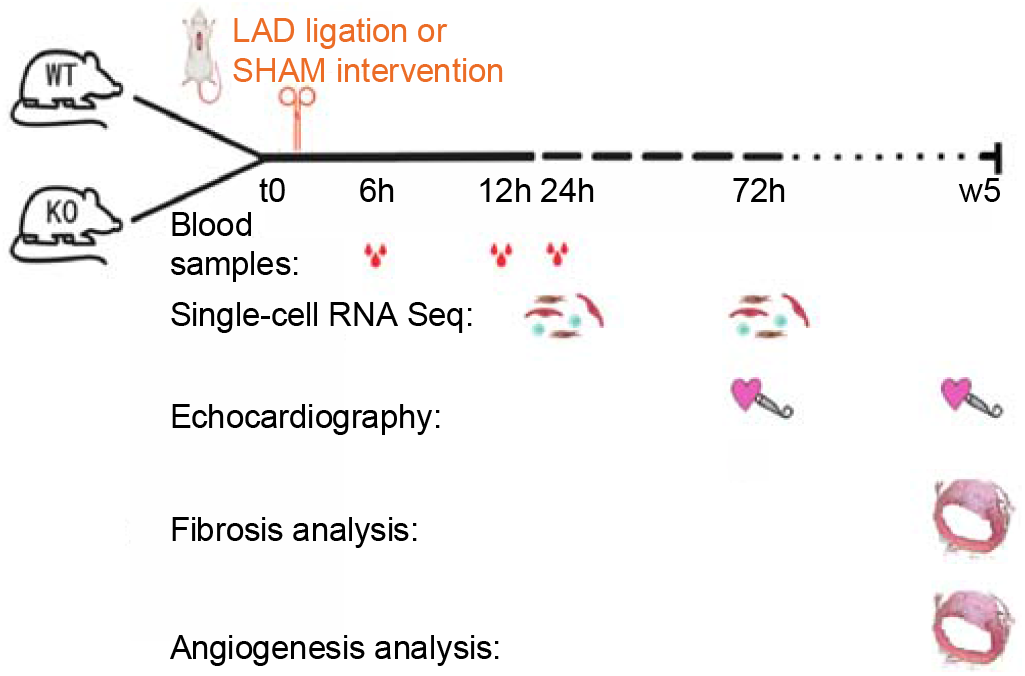
Overview of the study design including blood sampling, in-vivo echocardiography, single-cell RNA sequencing, and immunohistochemistry analysis of heart sections to assess fibrosis or angiogenesis using Col1a1 or CD31.

### Surgery

All mice included underwent surgery, which was performed by experimenters with multiple years of training in vascular microsurgery. The experimenters performing the surgery (sham or LAD ligation) were blinded for the genotype. The animals were anaesthetized by an intraperitoneal injection of a mixture of ketamine/xylazine/acepromazine (65, 15, and 2 mg/kg respectively), i.e. an injection volume of 0.1 mL per 10g of body weight. The skin was incised, the pectoral and intercostal muscles spread, to make an opening in the left third intercostal space. The pericardium was gently opened and a 7-0 suture was then passed under the left coronary artery approximately 1-2 mm below the tip of the left atrium. The ligature was then tightened, resulting in a blanching of the portion of the left ventricle supplied by the left coronary artery. The ligature was left in place (permanent ligation) and the animal’s rib cage closed (6-0 silk thread). A 5% glucose solution was administered intraperitoneally (0.3 to 0.5mL) and a local antiseptic (hydroalcoholic solution) was applied to the skin over the scar. Buprenorphine hydrochloride was administered as an analgesic after surgery as soon as the animal awakened. For sham animals, the same procedure was performed: the only difference was that the suture was not tied around the left coronary artery but only passed under it and removed.

### Echocardiography

The mouse was placed under anesthesia with isoflurane 4% in oxygen, 1L/min for induction, and then maintained under anesthesia via a nose cone using 1-1.5%, 1L/minute. While under anesthesia the following parameters were monitored: heart rate, body temperature (with rectal probe), ECG trace, and respiration. The echocardiography images were obtained using a dedicated machine of Visualsonic (Vevo2100) by placing warmed gel on the mouse. The average anesthetic period lasted 10 minutes. The data analysis was performed blindly, i.e. not knowing neither the genotype nor interventional group.

### Blood sampling

We used incision of the tail vein with a sharp blade until we obtained 100 ul of blood. Blood was left to clot at RT for one hour and then centrifuged at 1500 g for 15 min. The supernatant (=serum) was snap frozen.

### Non-myocyte cells isolation from sham-operated and infarcted mouse hearts

Non-myocytes were extracted as previously described^24^. In short, immediately after neck dislocation, the heart was extracted and rinsed in ice-cold HBSS. Thereafter, ventricles were cut in small (2×2 mm) pieces and incubated with thermolysin and papain for 30 min at 37°C under constant gentle shaking and resuspension every 10 minutes using a Pasteur pipette. After incubation the reaction was stopped using ice-cold HBSS and large debris were sorted out using a 300 μm mesh. Cardiomyocytes were discarded using low speed centrifugation (50 G) and a 40μm filter. Cells were subsequently incubated with calcein-AM (live) for 10 min at 37°C and DAPI was used to mark dead cells.

FACS was used to sort live cells. We used a fairly stringent gating (FSC Area - SSC Area, FSC Area - FSC Height, and SSC Area - SSC Height to exclude doublets. We used calcein AM and DAPI to select positive cells and discard dead cells. For each sample, we collected at least 100’000 cells per heart and pooled the sorted cells of 4 hearts.

### Single-cell RNA sequencing

A Chromium Next GEM Chip G (10X genomics) was loaded with 12’000 cells and the libraries prepared with the Chromium Single Cell 3L reagents v3.1 strictly following the manufacturer’s recommendations. Briefly, an emulsion encapsulating single cells, reverse transcription reagents and cell barcoding oligonucleotides was generated. After the actual reverse transcription step, the emulsion is broken and double stranded cDNA generated and amplified in a bulk reaction. This cDNA was fragmented, a P7 sequencing adaptor ligated, and a 3’ gene expression library generated by PCR amplification. Libraries were quantified by a fluorimetric method and their quality assessed on a Fragment Analyzer (Agilent Technologies). Cluster generation was performed with an equimolar pool from the resulting libraries using the Illumina Novaseq 6000 Cluster cartridge v1.5. Sequencing was performed on the Illumina Novaseq 6000 using SBS cartridge 100C v 1.5 kit reagents according to 10X Genomics recommendations (28 cycles read1, 10 cycles i7 and i5 index reads and 90 cycles read2).

### Processing of 10x Genomics Chromium single-cell RNA seq data

Fastq files were processed with cell ranger count with default settings (cellranger-6.0.0) on the mouse genome reference refdata-gex-mm10-2020-A. *filtered_feature_bc_matrix* (cells) data were used for QC analysis with Seurat (3.1.1) in R (R version 3.6.1 (2019-07-05)) using default settings. At data loading genes expressed in less than 10 cells were filtered. We kept cells with more than 1500 UMI counts and more than 600 detected features, as well as less than 30% mito RNA reads. Data was randomly subset to 3’100 cells per sample. After this step, the standard Seurat pipeline was followed for normalization, integration, scaling and PCA dimensionality reduction. We use the JackStraw function from Seurat to select the significant Principal Components (p<0.001, 81 PCs).

### Cluster and sub-cluster identification

We used the Seurat *FindNeighbors* method with dims = 1:81, annoy.metric = ‘cosine’ and k.param = 30. *FindClusters* was run at different resolutions. Based on UMAP representation and known gene markers annotation, the resolution 0.8 was chosen. This identified 27 clusters that were labeled according to their cell types. *FindSubCluster* was run on the T cells cluster. Sub clustering was also performed on: (1) cells from clusters labeled ‘Fibroblasts’ and ‘Epicardial cells’ and (2) cells from clusters labeled ‘Monocytes’ and ‘Macrophages’. The Seurat pipeline (scaling, PCA, UMAP, FindClusters) was re-run. Based on the JackStraw procedure, 44 PCs were kept for fibroblasts/epicardial cells, and 85 for monocytes/macrophages (p<0.001). For sub-clustering, we used resolution 0.6.

### Differential gene expression and enrichment analyis

We used Seurat *FindMarkers* (parameter ‘assay = RNA’) with statistical test MAST^25^ to identify differentially expressed genes between KO and WT in the different clusters and sub-clusters. We selected genes with an adjusted P values < 0.05 and an absolute fold-change >= 2. Gene ontology enrichment analysis was performed with *clusterProfiler*^26^.

### Differential proportion analysis (DPA)

We used source code from Farbehi et al. for DPA^27^. Null distributions were generated with parameters n=10000 and p=1 for the permutation procedure. *P* values were adjusted with Benjamini & Hochberg’s method.

### Proteomics analysis

Samples (sera) were measured on Evosep LC and timsTOF Pro mass spectrometer. The acquisition mode used was diaPASEF and samples were analysed with DIA-NN (available on Github: https://github.com/vdemichev/DiaNN) using default settings with match between runs as previously described^28^.

### Immunohistochemistry

We performed staining of Col1a1 and CD31 as previously described.^22^ In short, hearts of mice in the group D were frozen in OCT. Thereafter, we performed cryosections of 4μm every 100 μm. The sections were fixed using acetone and air dried. After quenching for endogenous peroxidase activity with BLOXALL and blocked further with goat serum slides were incubated with primary antibodies. Slides were subsequently washed with PBS and incubated with biotinylated antibodies, before incubating with VECTASTAIN ABC Reagent. After further washes, heart sections were incubated with AEC Substrate. The reaction was stopped by washing with distilled H_2_O. Lastly sections were counterstained with Harris Haematoxylin and slides were mounted.

## DATA AVAILABILITY

The raw and processed scRNA-seq data can be found under following accession number: GSE210159 (https://www.ncbi.nlm.nih.gov/geo/query/acc.cgi?acc=GSE210159). The results from the proteomics analyses are provided in supplementary data.

## RESULTS

### Knocking out *Trpm4* does not improve long-term survival five weeks after MI

We first monitored the survival of *Trpm4* WT and KO mice to assess the influence of the genotype on overall mortality after MI. While sham-operated WT mice showed no mortality, KO mice displayed a baseline mortality of 8.3%, which did not reach statistical difference. We observed a significantly increased mortality of 22.2% in WT mice after MI compared to sham, all of which happened within the first 3 days and most of which within the first 24 hours post-MI. The overall mortality in *Trpm4* KO mice of 21.2 % was almost identical to that of WT MI. However, the survival curve of KO MI only started to differ from that of KO sham after 3 weeks, even though this difference did not reach statistical significance. Furthermore, at week 3 post-MI survival continued to decrease in KO mice, whereas WT survival remained stable after day 3. We observed no significant difference in survival between both genotypes five weeks after MI. Thus, we conclude that knocking out *Trpm4* does not improve long-term survival five weeks after MI.

### Knocking out *Trpm4* does not improve echocardiography measurements of cardiac function after MI

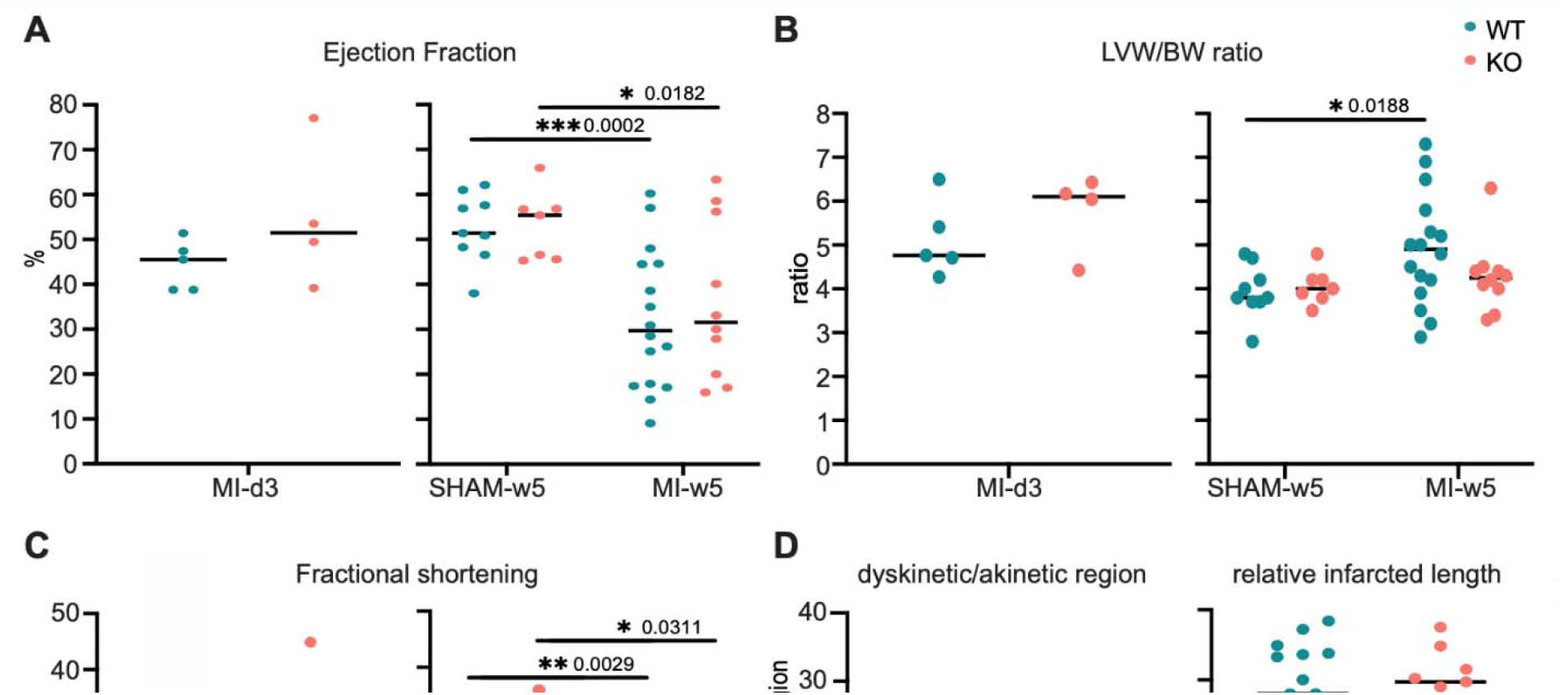

In order to assess cardiac function we used cardiac echography and confirmed that MI significantly reduced ejection fraction and fractional shortening as compared to sham in both genotypes. While ejection fraction remained preserved around 45-50% three days after MI, it decreased to approximately 30% in both genotypes five weeks after MI. We observed a similar phenomenon for the fractional shortening that was ∼20-25% three days post-MI, whereas it was down to approximately 15% five weeks after MI. Despite an initial trend towards an increased LVW/BW ratio in *Trpm4* KO mice compared to WT 3 days after MI, this trend was inversed 5 weeks after MI. This difference was not accounted for by changes in body weight as visualized in supplementary figure S1. Nevertheless, we observed no significant differences in ejection fraction, LVW/BW ratio, fractional shortening or dyskinetic region/relative infarct size between both genotypes after MI. A table including all values measured during echocardiography recordings is provided in the supplementary data. We conclude that knocking out *Trpm4* does not improve echocardiography measurements of cardiac function after MI.

### Serum proteomics changes in acute phase post-MI

In order to assess acute tissue injury and systemic inflammation we analyzed the sera of the mice at 6, 12, and 24h after surgery using LC-MS-based proteomics. As represented in Fig. 4B, the most highly detected protein in all our samples was albumin, which is the most abundant transport protein in the blood. Six hours after MI, we did not observe any significant differences between WT and KO sera (table provided in supplementary data). Twelve hours after MI, KO mice expressed significantly increased levels of pro-inflammatory acute phase protein Serum Amyloid P-Component (APCS) compared to WT. However, they displayed significantly lower levels of cardiomyocyte injury marker creatine kinase m (CKM), endothelial injury marker VE-cadherin (CDH5) as well as transmembrane receptor CD93, usually found on hematopoietic and endothelial cells, than WT. Moreover, the B-subunit of coagulation factor XIII (F13B) and transcobalamin II (TCN2) were also decreased in KO mice at this timepoint. Twenty-four hours after MI, CD93 remained lower in KO mice compared to WT, accompanied by surface glycoprotein CD44 and transmembrane receptor Ptprj, all typically expressed on hematopoietic cells. Their soluble form is a marker of injury and inflammation.^29–31^

**Figure 2.**
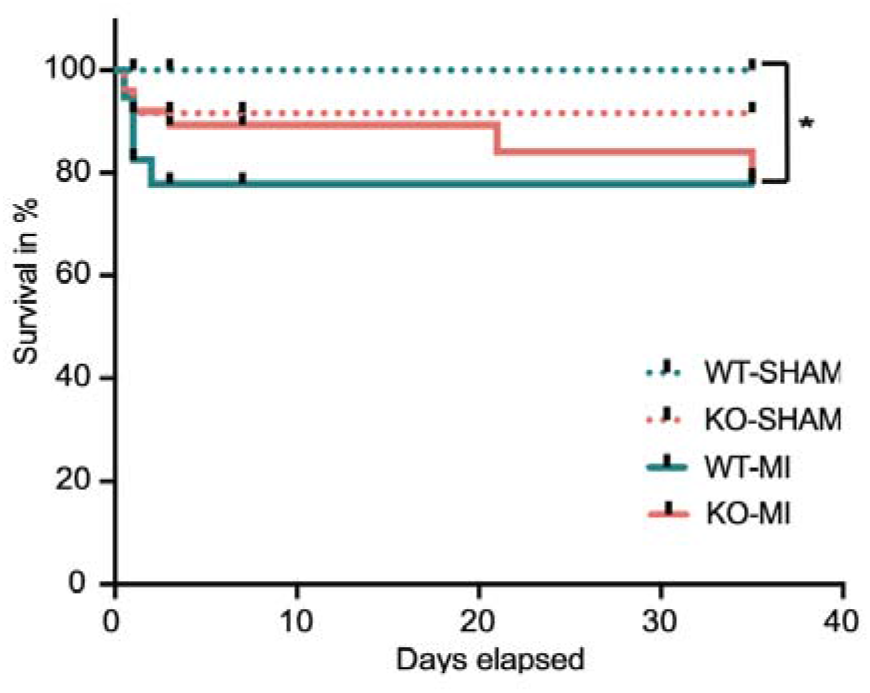
Survival curve for WT and Trpm4 KO mice during 5-week follow-up after LAD ligation. Every short black line represents a censored timepoint of sacrifice as described in the study design. Kaplan−Meier overall estimation was not significant using Gehan-Breslow-Wilcoxon test with P = 0.201. The only significant difference detected was between WT-SHAM and WT-MI with p = 0.0221 using Gehan-Breslow-Wilcoxon test, (n = 23 for WT-SHAM, 57 for WT-MI, 24 for KO SHAM, and 50 for KO-MI).

**Figure 4.**
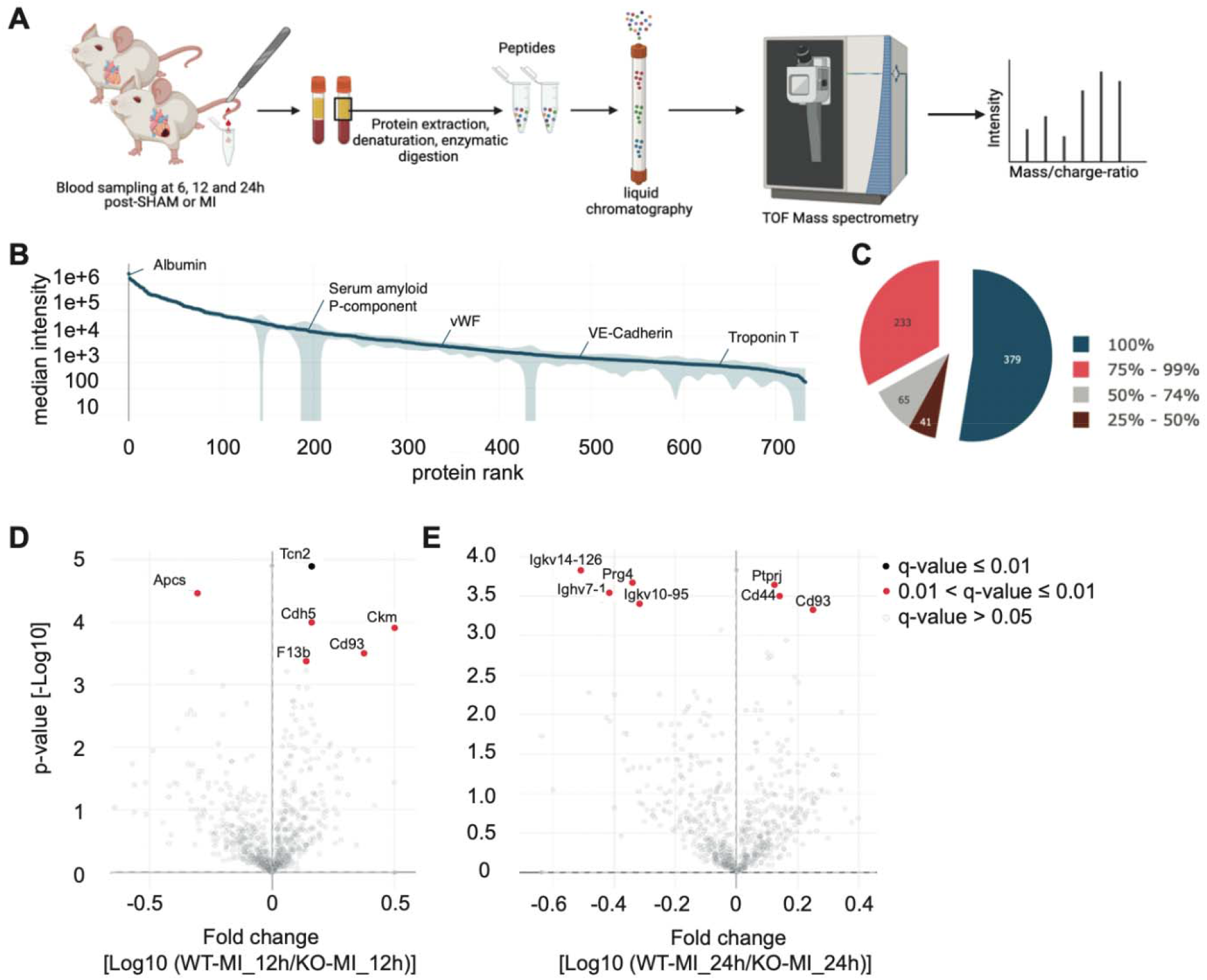
(A) Manual blood sampling and serum isolation followed by automated MS-based proteomics pipeline starting with 1 μl of serum, LC-MS equipment to generate MS raw data and data analysis. (B) In total, 718 proteins were quantified in this study, covering more than five orders of magnitude of MS signal. Examples of frequently applied biomarkers are labelled. (C) For 144 samples, 718 proteins were detected. The colour codes show the protein occurrences in % of samples, in which they were detected. (D) Volcano plot comparing the serum proteomes of 11 WT and 11 KO mice 12 hours after Ml. (E) Volcano plot comparing the serum proteomes of 6 WT and 13 KO mice 24 hours after Ml. The legend shows the colour code highlighting significant fold-depending on the q values.

On the other hand, *Trpm4* KO mice sera at 24 hours contained more immunoglobulin light chains such as Igkv14-126, Igkv10-95 and heavy chain Ighv7-1 as well as pro-reparative proteoglycan 4 (PRG4). Thus, these results are consistent with a stronger inflammatory process and lower tissue injury markers in KO compared to WT mice post-MI.

### MI induces more pro-inflammatory and pro-fibrotic cells in *Trpm4* KO than WT

Myocardial infarction triggers a complex immune response that involves many cellular and molecular players. In order to obtain the highest possible resolution into the acute inflammatory cellular response, we performed single-cell RNA sequencing 24h after sham or MI and 72h after MI. The cell types found in our non-myocytic population are shown in Fig. 5A, C and D. They comprised endothelial cell types 1, 2 and 3 (EC1, EC2, EC3 respectively), interferon inducible EC (i-EC), fibroblasts (fibro), smooth muscle cells (Sm), pericytes (Pc), granulocytes (Gc), interferon-inducible granulocytes (i-Gc), platelets (Tc), monocytes (Mono), macrophages (Mac), T and B lymphocytes (abbreviated T and B respectively) and dendritic cells (DC). Even though we used size filtering and centrifugation to exclude cardiomyocytes (Cm), we could observe a small contamination of the non-myocytic fraction. A deeper analysis of the monocytes/macrophages showed M1 monocytes (M1 Mono), M2 monocytes (M2 Mono), tissue-resident macrophages (TR-Mac), proliferating TR-Mac (pTR-M), M1 macrophages (M1), M2 macrophages (M2), pro-repair genes-high M2 macrophages (M2-Mac-R), interferon-inducible macrophages (i-Mac), Mac-6 macrophages (Mac-6), Mac-7 macrophages (Mac-7), monocyte-derived dendritic cells (MoDCs), and fibroblast-like Macrophages (Fibro-like Mac). The subcluster analysis of the fibroblast cluster showed proliferating fibroblasts (pFibro), epicardial cells (Epi), Sca1-high fibroblasts (F-SH), Sca1-low fibroblasts (F-SL), Wnt-expressing fibroblasts (F-Wntx), Myofibroblasts (Myo), and catalytic enzymes-high epicardial derived fibroblasts or myofibroblasts (F-phagocytic). Except for M2-Mac-R, which are not a distinct cell type, all these cell populations have been previously characterized^23,27,32–34^. A detailed UMAP visualization of exemplary marker genes per cell population, i.e. *Cdh5* for endothelial cells, *Acta2* for smooth muscle cells, *Kcnj8* for pericytes, *Col1a1* for fibroblasts, *Wt1* for epicardial cells, *Csf3r* for granulocytes, *Mafb* for monocytes and macrophages, *Cd209* for dendritic cells, *Cd3d* for T lymphocytes and Cd79a for B lymphocytes, can be found in the supplementary figure S3. 24 hours after MI, we measured significantly larger populations of granulocytes, monocytes and M1 macrophages in KO mice compared to WT. This correlated with an increased number of fibroblasts, myofibroblast and epicardial cells 72h after MI. Moreover, we detected the appearance of a population of fibroblast-like macrophages (fibro-like Mac), as well as a population of fibroblasts with an immune phenotype (F-phagocytic) 72h after MI in the KO, which were almost absent in WT mice. Overall, these results indicate that MI induces more pro-inflammatory, and more pro-fibrotic cells in *Trpm4* KO than WT mice.

**Figure 5.**
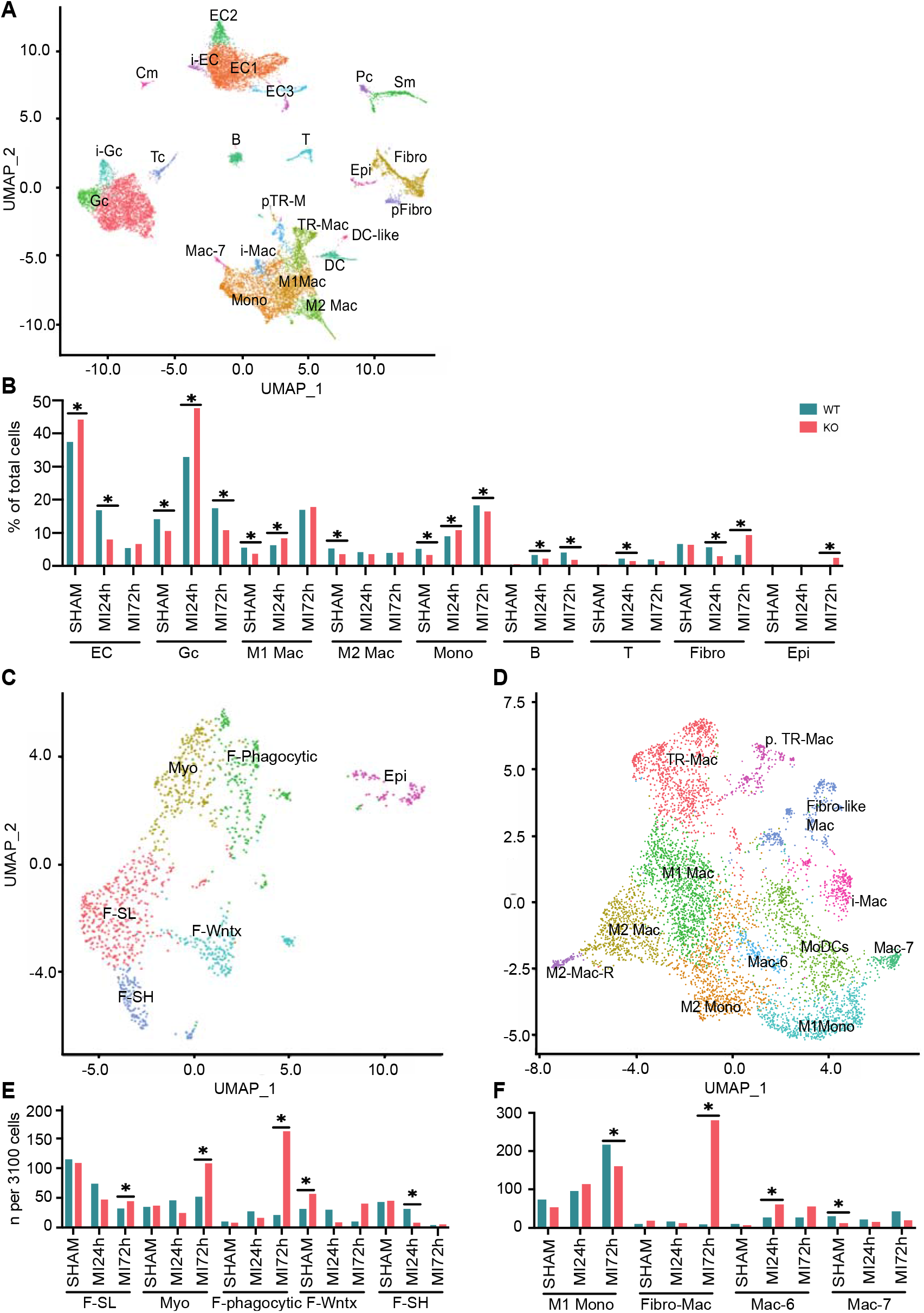
(A) UMAP plot showing detected lineages and sub-populations in NCM across conditions. (B) Cell population percentages across conditions with significant (* = adjusted p <0.05) differences between genotypes using Differential Proportion Analysis (DPA). (C) UMAP plot showing sub-cluster analysis after extracting fibroblasts/epicardial clusters from A. (D) UMAP plot showing sub-cluster analysis after extracting monocytes/macrophage clusters from A. (E) Cell population percentages of cells newly detected in C, analogous to B. (F) Cell population percentages of cells newly detected in D,

### *Trpm4* KO cells express higher levels of pro-inflammatory, -fibrotic and -angiogenic genes

In addition to the valuable and precise information about cell types that can be gained from single-cell RNA sequencing, we used pathway analyses and differential gene expression analyses to get a deeper understanding of the molecular processes at play. At 24h after MI endothelial cells started expressing higher levels of genes associated to wounding and acute inflammatory response, specifically response to cytokine Il1, leucocyte chemotaxis and adhesion, as well as coagulation. In line with this pro-inflammatory state, endothelial cells at 24h, at 72h, and M1 macrophages and fibroblasts at 72h showed positive regulation of peptide and protein secretion, as well as catalytic genes such as *Lyz2*. Moreover, genes associated with endopeptidases required for lysosomal processing and proteolysis were also increased. Furthermore, in KO endothelial cells and M1 macrophages genes promoting humoral and B-cell immunity were more highly expressed than in WT. This inflammatory response correlated with lower cellular movement, but increased angiogenesis-associated genes. In endothelial cells and fibroblasts, genes for divalent cation homeostasis, more precisely promoting Ca^2+^ mobilization, were increased 72h after MI compared to WT, but not in M1 nor M2 macrophages. Cell cycle analysis was done separately and can be seen in supplementary Fig. S4. 72h after MI, it consistently shows more TR-MAC, M1-Mac, M2-Mac, fibroblasts and epicardial cells in a proliferative state in KO compared to WT. In line with our subcluster analysis, the increased gene expression of cytokine *Il1b* and immune receptor *Fcer1g* in Fig. 6 exemplifies the increasing pro-inflammatory and immune phenotype of non-immune cells over time after MI in KO compared to WT.

**Figure 6.**
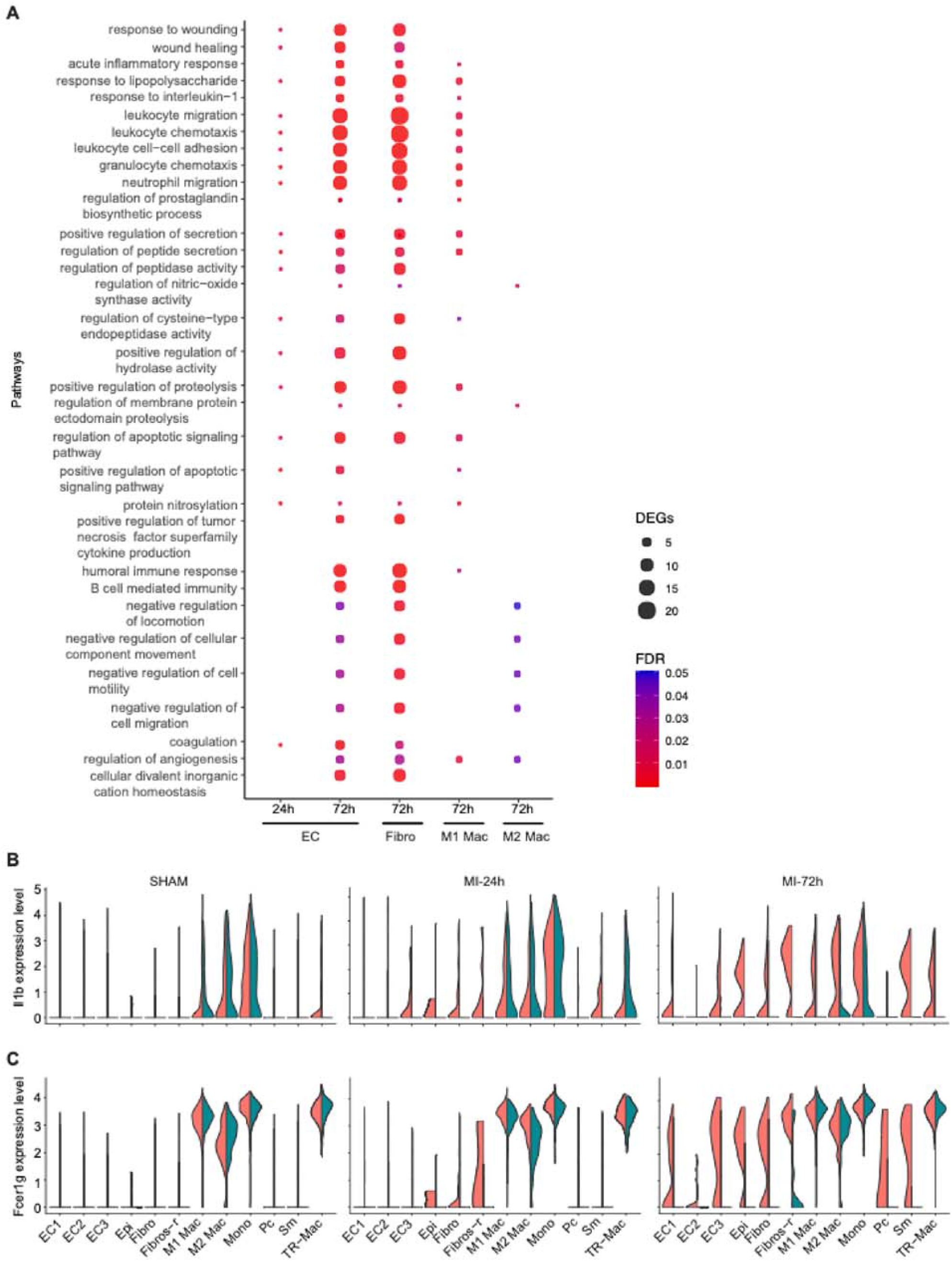
(A) Dot-plot visualizing gene ontology enrichment analysis. The top biological processes significantly enriched in at least two KO cell populations compared to WT are displayed. The size of the dots represent the number of differentially expressed genes (DEG) and the colour code shows the false discovery rate (FDR). (B) Violin plots show the differential gene expression of Il1b in cell populations, in which it was significantly up- or downregulated. (C) Analogous representation for gene

Taken together, these results indicate that MI induces a higher expression of pro-inflammatory, -fibrotic and -angiogenic genes in KO than WT mice. They further suggest that Ca^2+^ homeostasis is upregulated in KO compared to WT.

### *Trpm4* KO increases fibrosis and angiogenesis post-MI

In order to assess the long-term consequences of the cell types and pathways affected during acute inflammation, we performed immunostainings on hearts 5 weeks after surgery to quantify fibrosis. Fig. 7A shows the absence of scar tissue after sham surgery in both genotypes. After MI, however, large fibrotic scars are visible. The fibrotic area, including scar tissue and remote fibrosis, extends further in KO compared to WT hearts. The quantification of Col1a1-positive staining from at least 10 sections per heart from at least 6 hearts per experimental group as displayed in Fig. 7B shows that while MI leads to fibrosis in both genotypes, post-MI fibrosis was significantly higher in KO mice compared to WT. To assess angiogenesis, we used anti-CD31 to stain endothelium. Representative images of the heart slices are provided in supplementary figure S9. The quantification from Fig. 7C shows that KO mice displayed significantly more endothelial staining than WT mice after MI. In line with genes that we found upregulated acutely after MI, these results suggest that *Trpm4* KO promotes fibrosis and angiogenesis post-MI.

**Figure 7.**
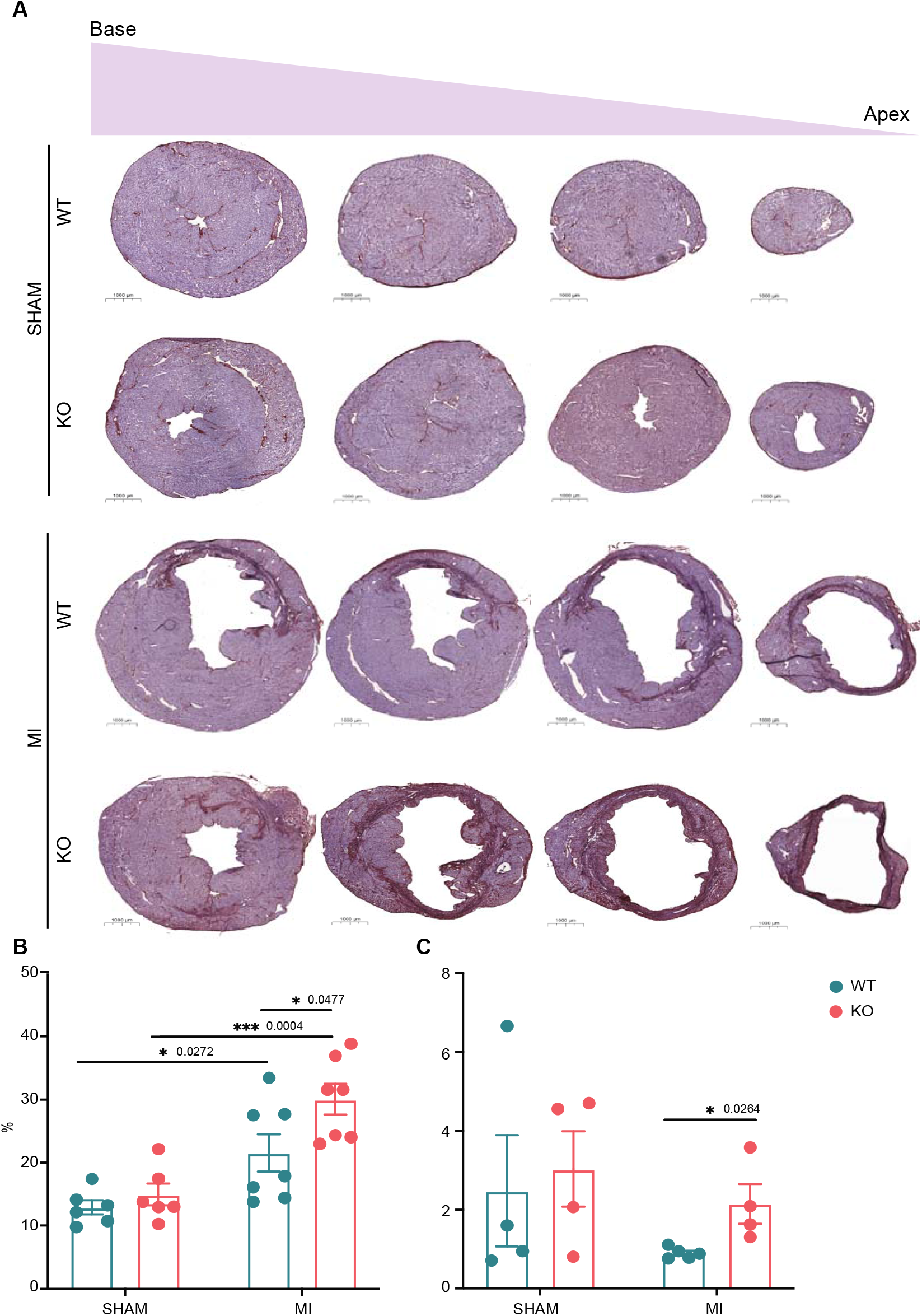
(A) representative images of transverse heart slices stained with Hematoxylin in purple and immunostaining of Col1a1 in brown. (B) Quantification of fibrosis by calculating Col1a1-positive surface as a percentage of total HE-positive surface. (C) Analogous quantification of angiogenesis using CD31 as endothelial marker instead of Col1a1.

## DISCUSSION

This study represents the first investigation on the role of TRPM4 in the global inflammatory and healing process following myocardial infarction in mice. The main observations were that absence of TRPM4 increases acute inflammatory molecular and cellular markers, cardiac fibrosis, and angiogenesis.

Over the course of five weeks, we did not find significant differences in survival after MI between both genotypes, even though we included more mice than any other study of this type. This correlated with our functional data showing no differences in echocardiography parameters at week 5, which is also in line with the baseline echocardiography measurements of Jacobs et al. after ten week^13^.

The trend showing a decreased early mortality in KO correlated with significantly lower tissue injury markers such as muscle-specific creatine kinase m (CKM)^35,36^ and endothelial-specific VE-Cadherin (CDH5)^37–40^ in the serum proteomics analysis 12h after MI. This is in agreement with the findings of Piao et al. showing that *Trpm4* gene deletion and pharmacological inhibition protect cardiomyocytic H9c2 cells from cell death upon ischemia-reperfusion injury as mimicked by H_2_O_2_ incubation^5^. Similarly, Becerra et al. found that silencing or pharmacological blocking of TRPM4 in human umbilical vein endothelial cells significantly reduce lipopolysaccharide-induced cell death^41^.

At the same time, KO was associated with a stronger acute inflammatory phenotype, showing higher levels of the pentameric serum amyloid P component (APCS). APCS is a sensitive marker of inflammation in mice, analogous to CRP in humans^42^. It recognizes and opsonizes necrotic, as well as apoptotic cells and mediates their phagocytosis through type I and type III Fc-receptor gamma, also called Fcer1g and Fcer3g respectively^43^. Thus, higher levels of APCS should result in increased opsonization, faster phagocytosis and quicker resolution of inflammation. Moreover, APCS has been shown to inhibit maladaptive processes such as fibrosis, fibroblast proliferation and monocyte to fibrocyte transition^43–45^. Contrarily to this hypothesis, the *Trpm4* KO mice, which had higher APCS levels also showed increased fibrosis, more fibroblast proliferation and displayed a fibrocyte-like monocytic population at day 3 after MI, which was almost absent in WT mice. Since the above-mentioned effects of APCS are mediated by Fcer1g on monocytes, and imply a proper monocytic function, we hypothesized that an impaired phagocytic function in KO monocytes and macrophages may explain this unexpected observation. Indeed, a study performed by Serafini et al. demonstrated that deletion of *Trpm4* impairs phagocytic function of macrophages during sepsis in mice.^17^. They observed that impaired Ca^2+^ mobilization alters monocytes’ and macrophages’ phagocytic activity in KO, leading to increased pro-inflammatory Ly6C^+^ monocytes and pro-inflammatory cytokine production. Analogous to their results, we found *Trpm4* KO to be associated with upregulated genes responsible for cellular divalent inorganic cation homeostasis, stronger recruitment of pro-inflammatory neutrophils, M1 monocytes and macrophages, as well as stronger expression of pro-inflammatory genes such as *Il1b, Ly6c* (also called *Lyz2*), *S100a8, S100a9, Fcer1g, Cd74, Cd52, C1qb* and *Tyrobp*.

Furthermore, this increased inflammation in *Trpm4* KO mice was associated with marked fibroblast proliferation, and a larger myofibroblast population at day 3 post-MI. In turn, this pro-fibrotic signature correlated with increased fibrosis as measured by type I collagen staining at five weeks post MI. Indeed, it is well established that overshooting inflammation is a driver of cardiac fibrosis after MI^22,32,46–48^. The increased fibrosis we observe in KO mice does not support the hypothesis of Simard et al. that TRPM4 drives cardiac fibrosis^49^. However, their assumption was based on their *in vitro* measurement of slower fibroblast growth and decreased transition into myofibroblasts upon TRPM4 inhibition. It can be suggested that in the context of MI, this effect is overridden by the role of TRPM4 in inflammation.

Considering what is known about TRPM4 on calcium homeostasis in endothelial cells and fibroblasts, our findings support the hypothesis that knocking-out *Trpm4* promotes calcium-influx. Indeed, we found the pathway that upregulates divalent inorganic cation homeostasis increased in KO endothelial cells and fibroblasts compared to WT. Consistent with the findings of Serafini et al. showing that the physiological effect of *Trpm4* KO on calcium influx in murine macrophages is abolished when macrophages are exposed to an inflammatory environment, we did not observe this pathway upregulated in M1 or M2 macrophages.

Along with the above-mentioned pro-inflammatory genes, our differential gene and pathway analysis also showed increased pro-angiogenic signatures in KO cardiac cells 3 days after MI. This is consistent with increased angiogenesis measured by CD31 staining five weeks post-MI, and in line with extensive previous research showing that inflammation is a key promotor of angiogenesis after MI^50^. In addition, *Trpm4* silencing or inhibition has also been shown to promote angiogenesis in stroke^51^. Interestingly, there is one protein that has been associated to most of our observations: Factor XIIIa, the last protein of the coagulation cascade that is also crucial for wound healing and adequate cardiac remodeling after MI. It does so by promoting inflammation, increasing neutrophil recruitment, enhancing angiogenesis and Col1a1 synthesis, as well as upregulating fibroblast proliferation. FXIIIa-depleted mice die of cardiac rupture with a 5-days mortality of 100% ^52–55^. Furthermore, our sera analysis showed elevated factor XIIIb 12h after MI in WT compared to KO. FXIII is a tetramer consisting of two A- and two B-subunits non-covalently bound. They are also called FXIIIa and FXIIIb respectively. Once activated by thrombin, Factor XIII releases its two A-subunits to stabilize the fibrin clot. While the A-subunits are mainly produced in monocytes, macrophages and megakaryocytes/platelets, the B-subunits are synthesized only in hepatocytes. The main function of FXIIIb is to transport the hydrophobic A-subunits within the plasma. Accordingly, all circulating FXIIIa in the plasma is bound to FXIIIb, whereas FXIIIb is present in excess and only half of it is bound to FXIIIa^53^. Since FXIII is not an acute phase protein^55^, the increased FXIIIb we observe in WT 12h after MI is unlikely caused by increased synthesis. It seems much more likely to depict a relative increase in its unbound form, which would result from lower FXIIIa levels. Indeed, our single-cell data suggested that proliferating tissue-resident macrophages and fibroblasts expressed less FXIIIa in WT than KO after MI supporting this hypothesis. In line with this, our gene ontology analysis confirmed less genes associated with coagulation cascade in WT compared to KO after MI. Thus, the proteomic analysis, the differential gene analysis as well as the pathway analysis would support the hypothesis that MI induces lower levels of FXIIIa in WT compared to KO. Interestingly, FXIIIa is also a transglutaminase. Noll et al. showed that its deposition at endothelial cell junctions decreases endothelial permeability in vitro as well as in vivo after myocardial infarction in rats^56^. Similarly, we also observed increased soluble VE-Cadherin in the sera of the WT mice, which is a marker of endothelial barrier disruption and increased permeability upon inflammation^57,58^. This suggests that *Trpm4* KO may decrease endothelial permeability after myocardial infarction as has been shown in the brain upon stroke^59,60^.

One obvious limitation of this study is that it investigates the role of Trpm4 in mice from a C57Bl/6N strain, while Medert et al. demonstrated that these mice respond differently to MI than mice from a 129SvJ background^20^. Thus, the latter may display a different inflammatory response upon MI. Despite the fact that we included a sample size, which is unprecedently high for any study investigating TRPM4 in MI to this date, the survival analysis we performed did not show any genotype-dependent difference in mortality, contrarily to what has been observed by Jacobs et al.^13^ In line with their findings, and with the present results showing significantly decreased tissue injury markers in KO, we found a trend towards a lower first-week mortality in *Trpm4* KO. However, in order to test whether this difference in early survival of about 11% is significant, we would need a sample size of at least 178 mice per group. Inflicting such a high severity grade intervention on such a large number of mice would defy the 3R (replace, reduce, refine) principles for animal welfare protection, which is why we did not attempt to increase the sample size. Another limitation of this study is that we did not include female mice for the same 3R reasons. However, the key findings of the present study warrant to be replicated in female mice in the future. Furthermore, since we performed our analysis on serum, we could not measure FXIIIa, as it can only be assessed in plasma ^61^. Moreover, we used a global KO strategy. Hence, our study offers promising target proteins and cells involved in the pro-inflammatory phenotype of *Trpm4* KO, but it is impossible to decipher with certainty, via which cells and mechanisms this happens. For that purpose, studies including an inducible knockout or a specific TRPM4 inhibitor, as well as cell-type-specific knock-outs will be necessary.

Taken together, our data show that knocking out *Trpm4* does not confer any long-term survival benefit or improved cardiac function over the course of five weeks. Nevertheless, *Trpm4* KO promotes recruitment of inflammatory cells to the heart and upregulates inflammatory genes in macrophages, endothelial cells as well as fibroblasts. Moreover, it stimulates proliferation of macrophages and fibroblasts 72h after MI, which correlates with increased fibrosis and angiogenesis five weeks post-MI. Thus, we demonstrate that TRPM4 plays a two-sided role in MI. On the one hand increased fibrosis promotes ventricular stiffness and heart failure, while on the other hand increased angiogenesis is associated with improved cardiac recovery post-MI. Therefore, its inhibition as has been proposed by many previous studies, bears risks. It may be indicated to carefully time and target its inhibition.

## Supporting information

Gene Ontology

differential prop. analysis clusters

differential prop. analysis subclusters

diff. gene analysis clusters

differential prop. analysis cell cycle

Echos 72h

echos 5 weeks

proteomics 6h

proteomics 12h

proteomics 24h

proteomics raw

## ACKNOWLEDGMENTS

We thank

- Anne Catherine Clerc, Dr. Corinne Berthonneche and Dr. Alexandre Sarre from the CAF Facility, University of Lausanne/CHUV
- From the GTF Facility University of Lausanne: Julien Marquis, Karolina Bojkowska and Corinne Peter
- Omicera GmbH
- Dr. Mohamed Nemir, from the University of Lausanne for his help on immunostainings
- Danny Labes from the FACS Facility at the University of Lausanne
- Dr. Stefan Müller from the FACS Facility at the University of Bern
- Anastasia Milusev, from the DBMR, university of Bern, for her readiness to share material
- Figures 2 and 4 were created with biorender.com

## ABBREVIATIONS

MI: Myocardial infarction
WT: wildtype
KO: Knock-out

