## Supplementary Figures for "Deletion of the ion channel *Trpm4* increases cardiac inflammatory markers and fibrosis after myocardial infarction in mice"

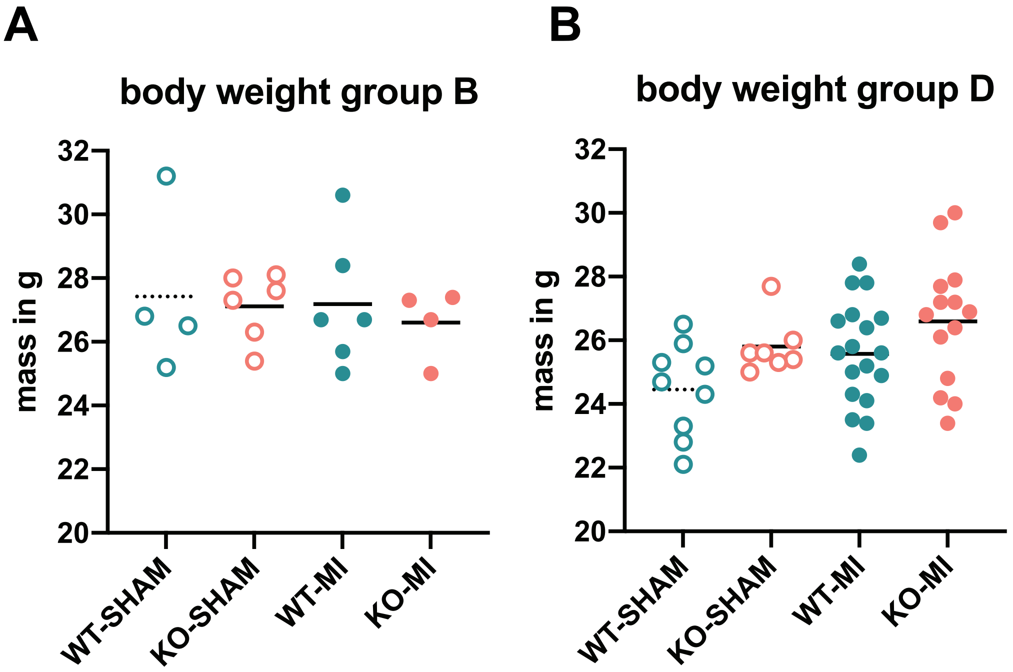


C

| | P-vaule |
|---|---|
| WT-SHAM vs. KO-SHAM | 0.0517 |
| WT-MI vs. KO-MI | 0.1226 |
| WT-SHAM vs. WT-MI | 0.0909 |
| KO-SHAM vs. KO-MI | 0.2225 |

**Figure S1.** (A) shows the body weights in gramms of the mice in group B at the timepoint of sacrifice at 72h. (B) shows the body weights in gramms of the mice in group D at the timepoint of sacrifice at 5 weeks. (C) The table shows the comparison between the different conditions of group D displayed in fig. S1B.

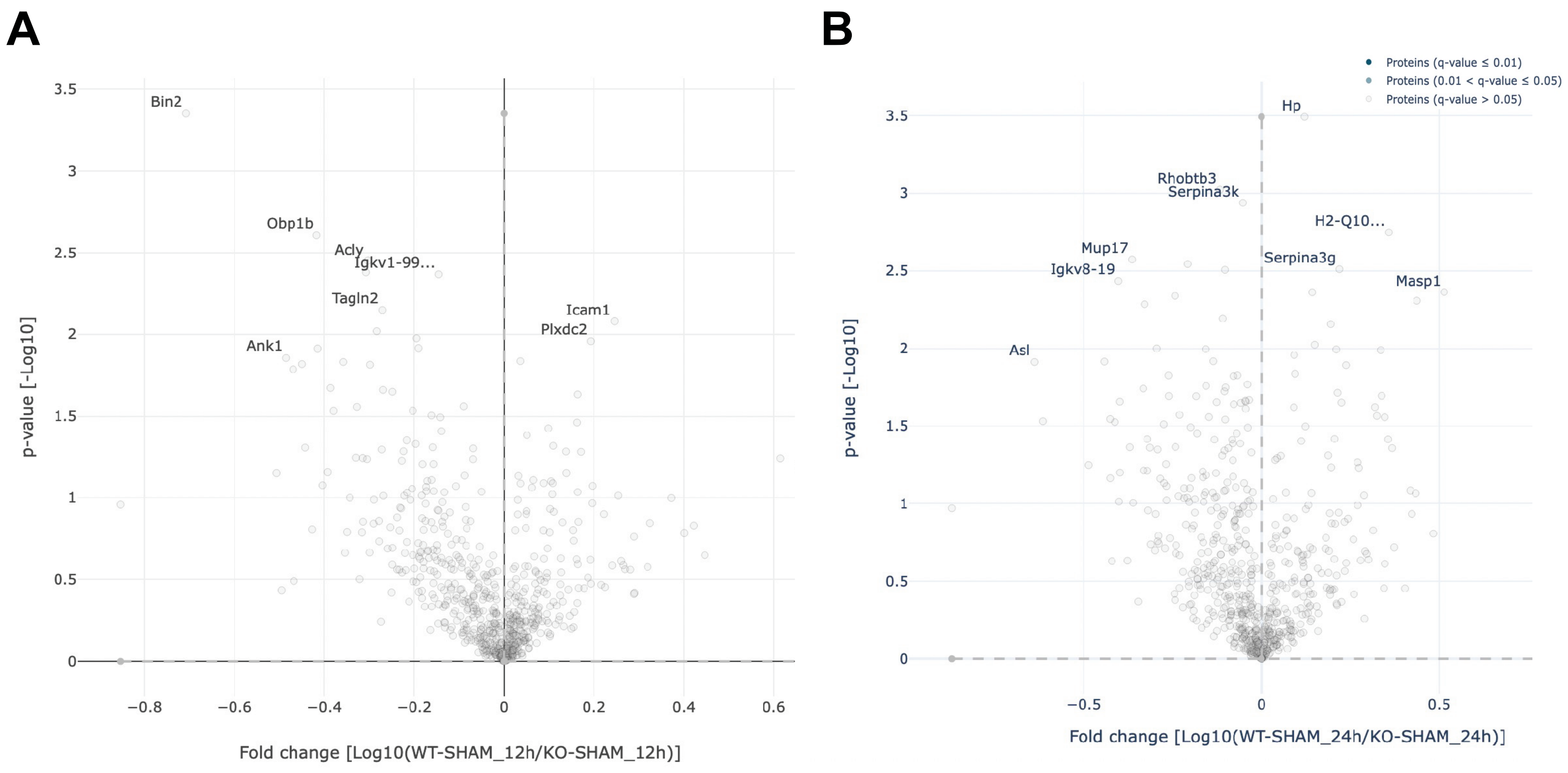


**Figure S2.** (A) Volcano plot comparing the serum proteomes of 6 WT and 5 KO mice 12 hours after SHAM surgery. (B) Volcano plot comparing the serum proteomes of 7 WT and 4 KO mice 24 hours after SHAM intervention. The legend shows the colour code highlighting significant fold-depending on the q values.

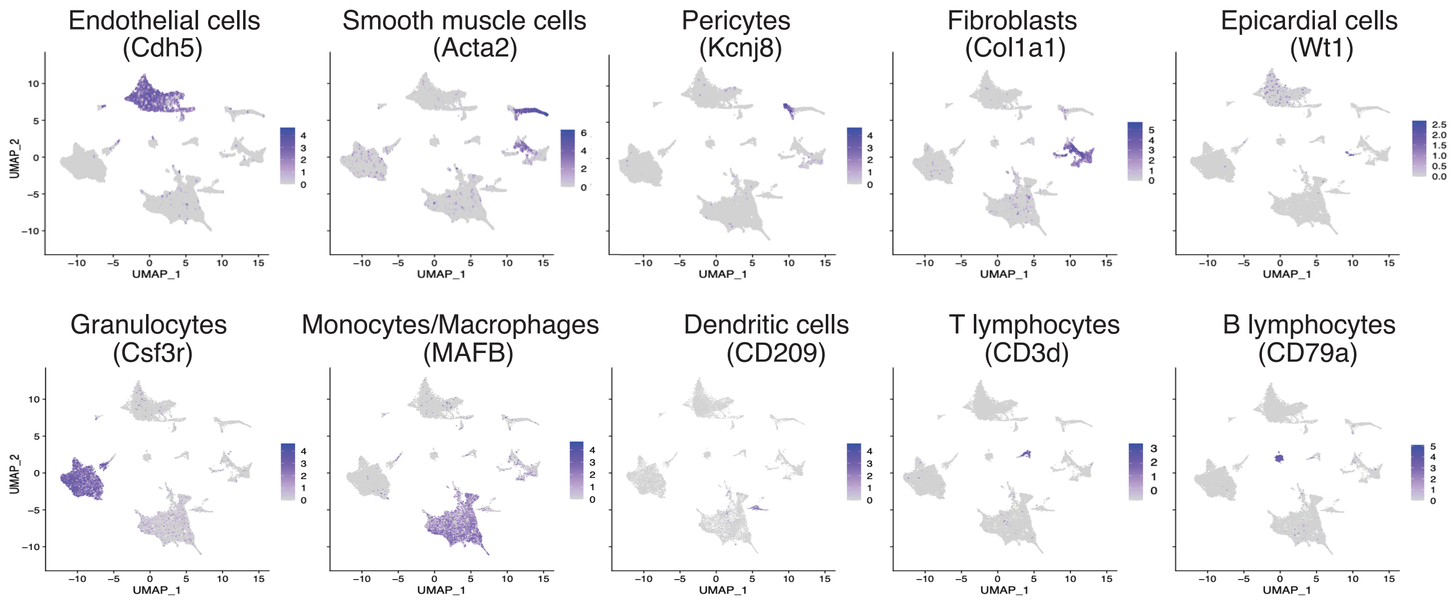


**Figure S3.** UMAP visualization of all integrated samples, with gene expression of cell type markers labeled in brackets colour coded in purple.

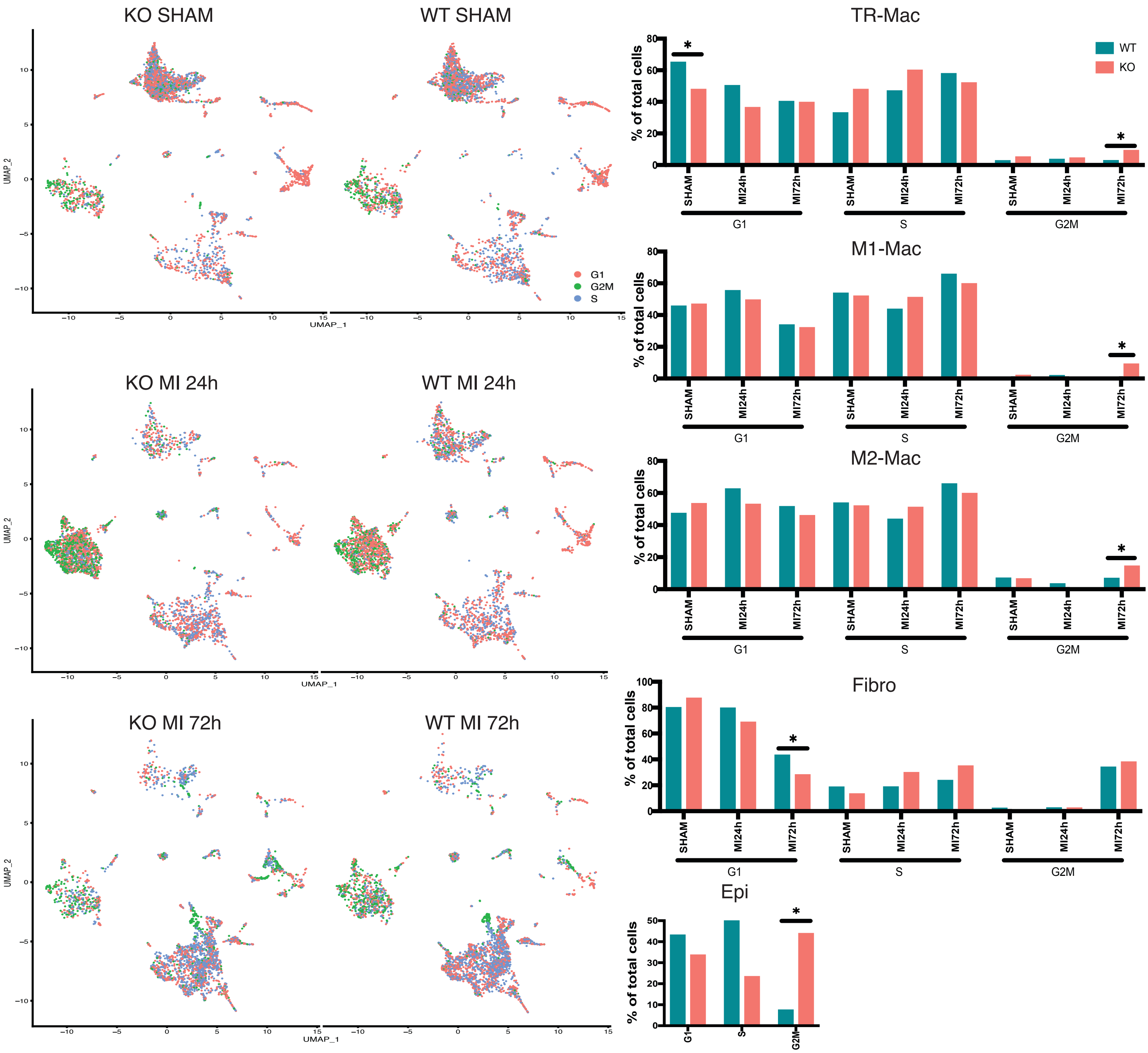


**Figure S4 .** (A) UMAP visualization of cell cycle stage according to colour code 24h after SHAM intervention in WT or KO; (B) same visualization for samples 24h after MI in WT or KO (C) same visualization for samples 72h afer MI in WT or KO (D) Percentage of cells per cell cycle stage and cell type. Cell population percentages across conditions with significant (* = p <0.05) differences between genotypes using Differential Proportion Analysis (DPA) and p-value adjustment after multiple testing correction.

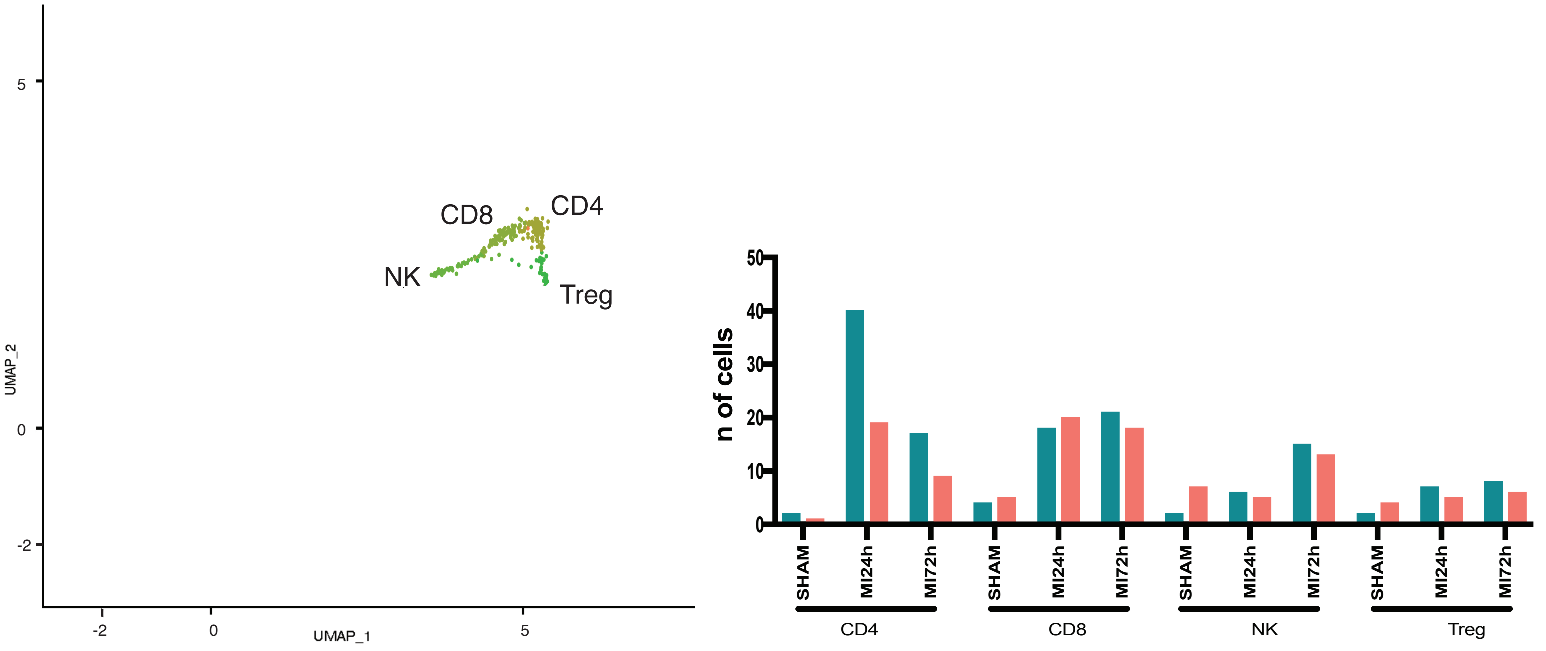


**Figure S5.** (A) UMAP plot showing sub-cluster analysis after extracting Tcell cluster from fig.5A; (B) Cell population percentages across conditions with significant (* = p <0.05) differences between genotypes using Differential Proportion Analysis (DPA) and p-value adjustment after multiple testing correction.

**Figure S6.** Heatmap of differentially expressed genes for macrophages/monocytes sub-cluster.

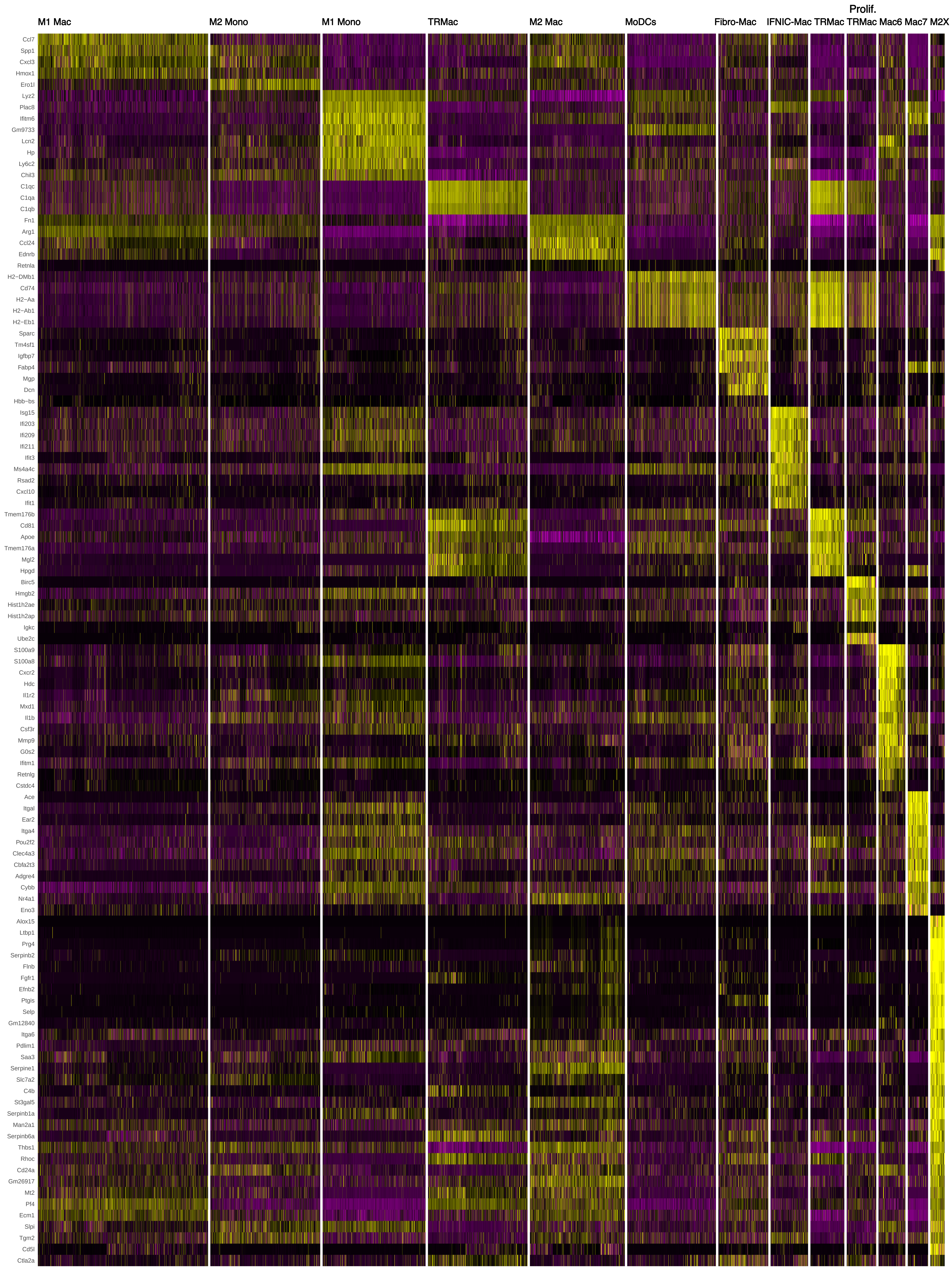

**Figure S7.** Heatmap of differentially expressed genes forr Fibroblasts sub-cluster.

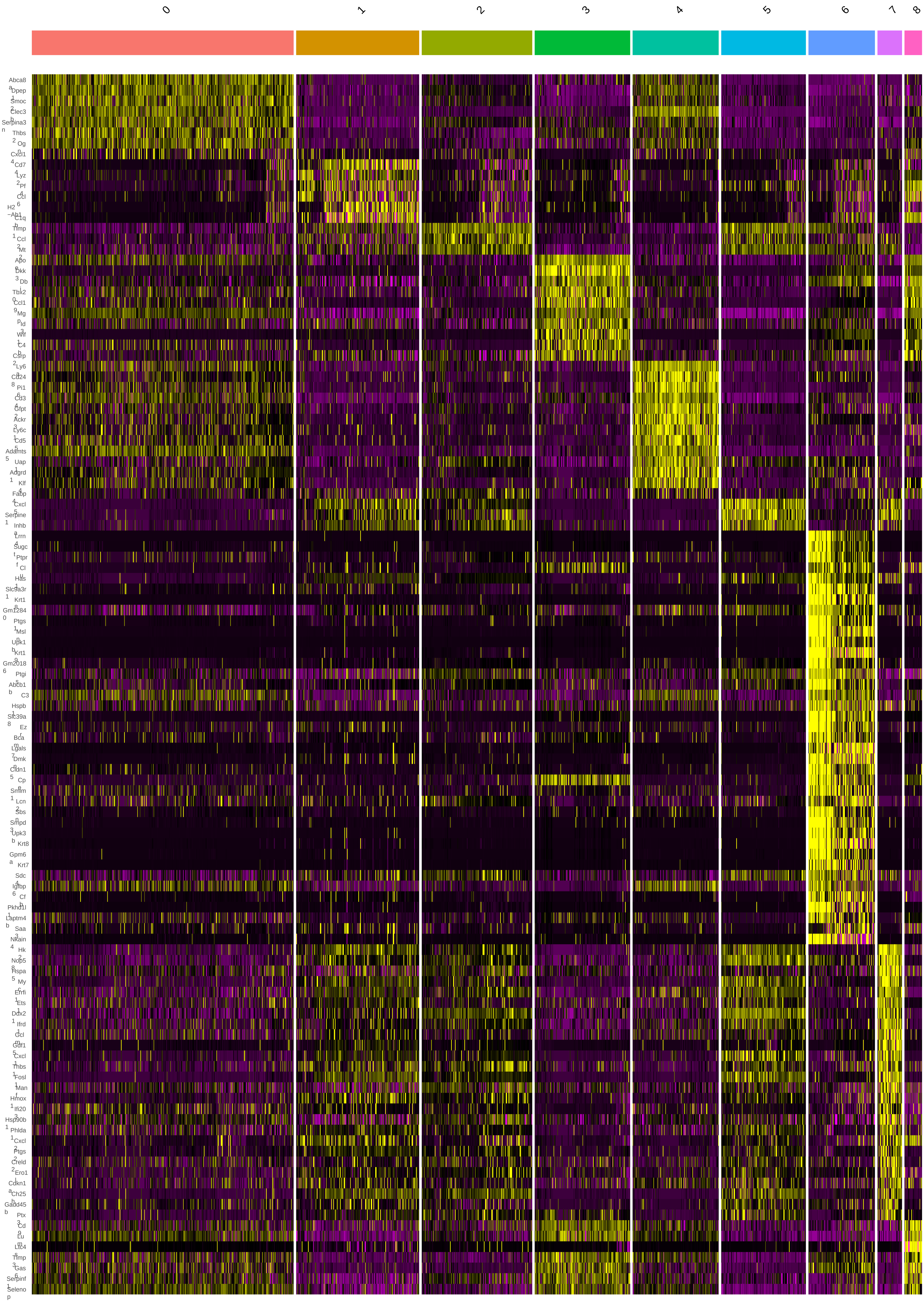

**A**

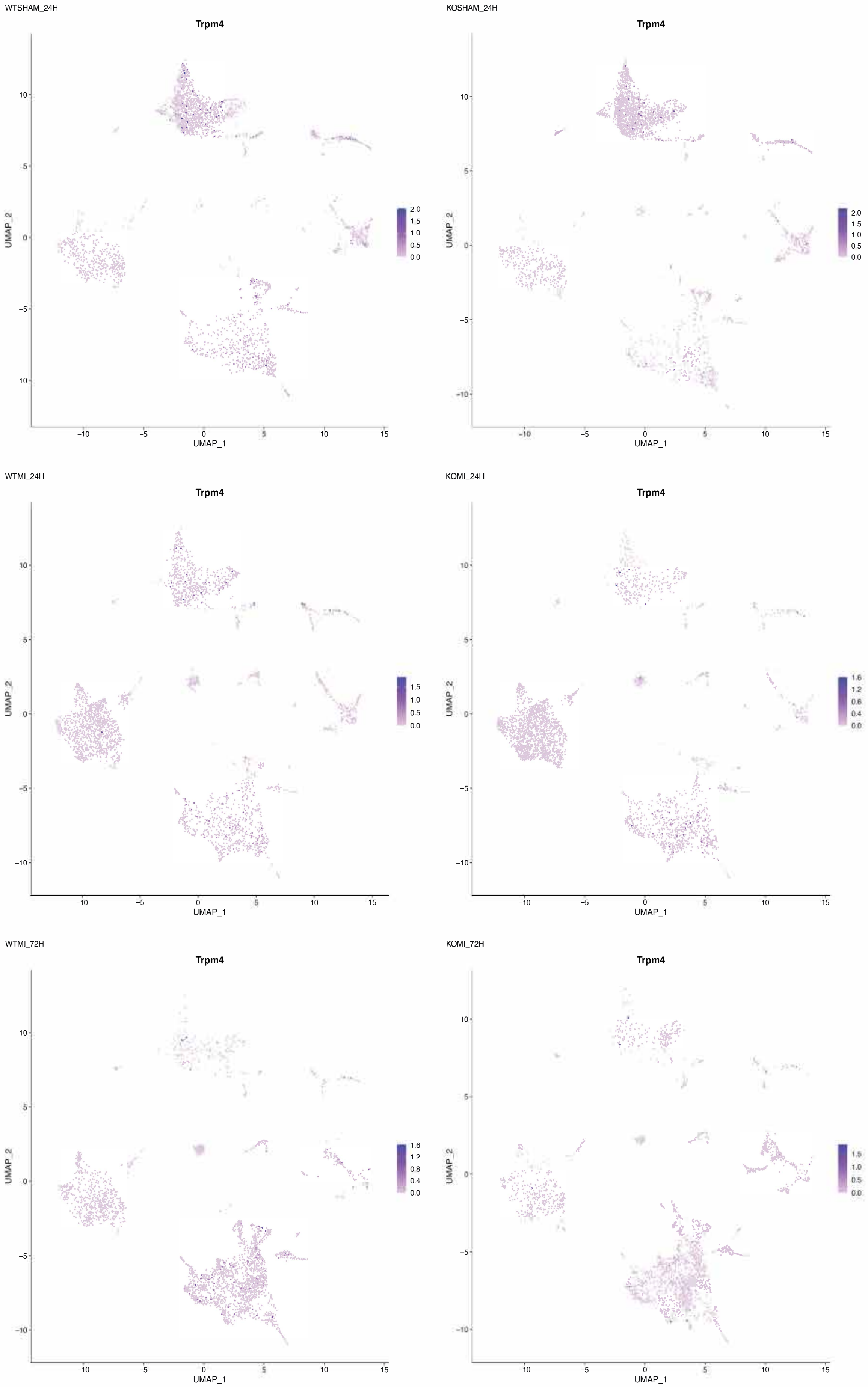


**B**

| | Ratio TRPM4 to PolR2A (2^-DeltaCq) |
|---|---|
| WTSHAM | 1.086039897 |
| WTMI24h | 0.509133636 |
| WTMI72h | 0.314219691 |

**Figure S8.** (A) UMAP plots showing relative TRPM4 expression per condition. (B) gene expression ratio of TRPM4 relative to Polr2a measured by qPCR on total non-myocytic cells from same samples as in A.

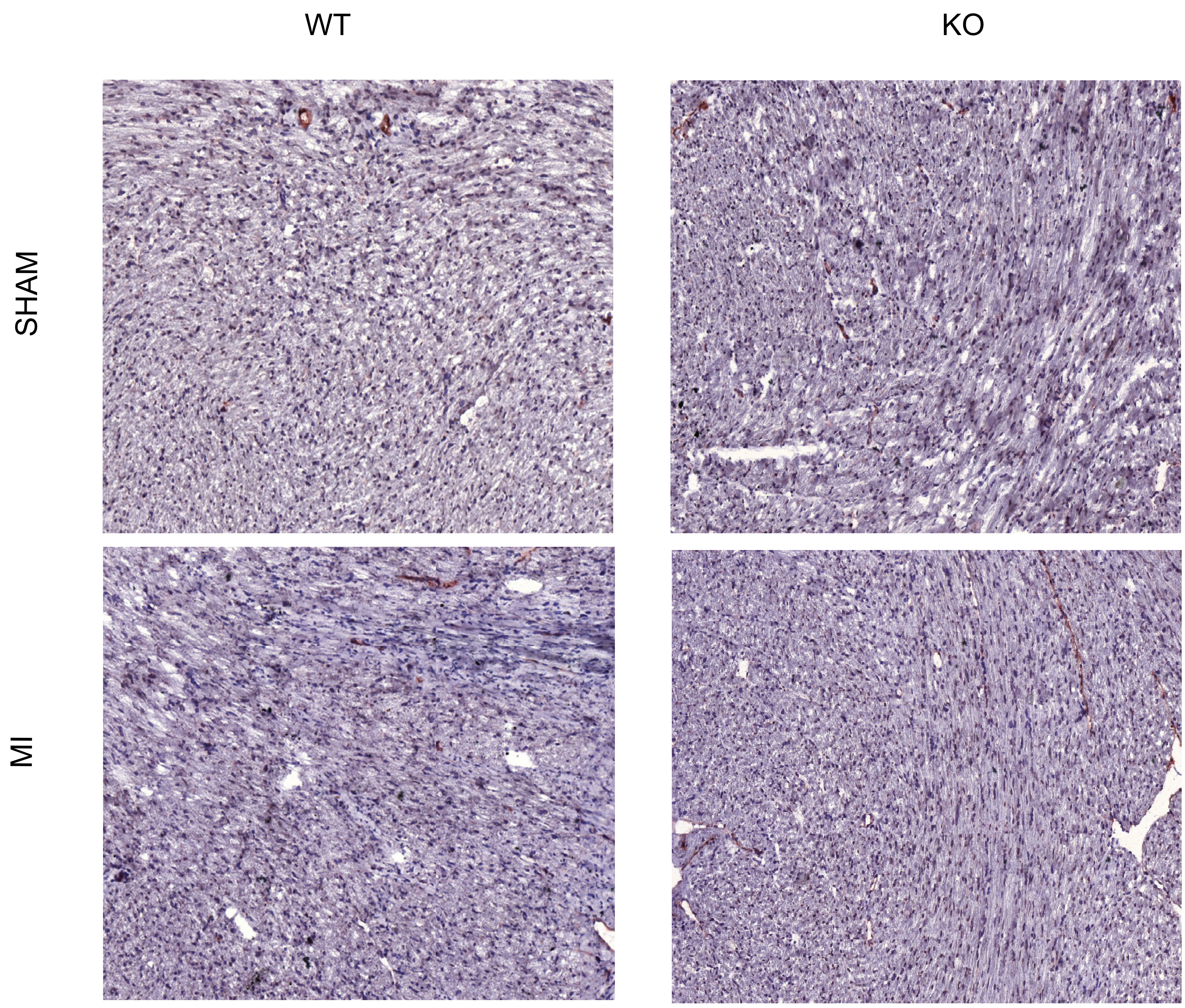


**Figure S9.** Representative images of CD31 immunostaining co-stained with HE five weeks after SHAM or MI surgery in WT or KO mice.
